# The selenophosphate synthetase, *selD*, is important for *Clostridioides difficile* physiology

**DOI:** 10.1101/2021.01.06.425661

**Authors:** Kathleen N. McAllister, Andrea Martinez Aguirre, Joseph A. Sorg

**Affiliations:** Department of Biology, Texas A&M University, College Station, TX; Skirball Institute of Biomolecular Medicine, New York University School of Medicine, New York, NY

## Abstract

The endospore-forming pathogen, *Clostridioides difficile*, is the leading cause of antibiotic-associated diarrhea and is a significant burden on the community and healthcare. *C. difficile*, like all forms of life, incorporates selenium into proteins through a selenocysteine synthesis pathway. The known selenoproteins in *C. difficile* are involved in a metabolic process that uses amino acids as the sole carbon and nitrogen source (Stickland metabolism). The Stickland metabolic pathway requires the use of two selenium-containing reductases. In this study, we built upon our initial characterization of the CRISPR-Cas9-generated *selD* mutant by creating a CRISPR-Cas9-mediated restoration of the *selD* gene at the native locus. Here, we use these CRISPR-generated strains to analyze the importance of selenium-containing proteins on *C. difficile* physiology. SelD is the first enzyme in the pathway for selenoprotein synthesis and we found that multiple aspects of *C. difficile* physiology were affected (*e.g*., growth, sporulation, and outgrowth of a vegetative cell post-spore germination). Using RNAseq, we identified multiple candidate genes which likely aid the cell in overcoming the global loss of selenoproteins to grow in medium which is favorable for using Stickland metabolism. Our results suggest that the absence of selenophosphate (*i.e*., selenoprotein synthesis) leads to alterations to *C. difficile* physiology so that NAD^+^ can be regenerated by other pathways.

**Importance:** *C. difficile* is a Gram-positive, anaerobic gut pathogen which infects thousands of individuals each year. In order to stop the *C. difficile* lifecycle, other non-antibiotic treatment options are in urgent need of development. Towards this goal, we find that a metabolic process used by only a small fraction of the microbiota is important for *C. difficile* physiology – Stickland metabolism. Here, we use our CRISPR-Cas9 system to ‘knock in’ a copy of the *selD* gene into the deletion strain to restore *selD* at its native locus. Our findings support the hypothesis that selenium-containing proteins are important for several aspects of *C. difficile* physiology – from vegetative growth to spore formation and outgrowth post-germination.

## Introduction

*Clostridioides difficile* is a major concern as a nosocomial and community-acquired gut pathogen (1). This pathogen has become the most common cause of health care-associated infections in United States hospitals (2, 3). In 2017 in the United States, approximately 223,900 cases of *C. difficile* infection were identified and nearly 12,800 of those resulted in death. In the same year, it was estimated that *C. difficile* infections resulted in more than $1 billion in excess health-care costs (4). The burden of this pathogen on patients, the community, and health-care has led the Centers for Disease Control and Prevention to identify this bacterium as an urgent threat (5).

Key to battling this aggressive pathogen is understanding the basic processes *C. difficile* uses to complete its lifecycle. While our understanding of *C. difficile* physiology has increased dramatically in the last decade, specifically in toxin production / regulation, sporulation and germination, our understanding of metabolic processes is lacking (6–11). A recent review by Neumann-Schaal *et al*. nicely discussed the known metabolic processes involved in energy generation in *C. difficile* (12). *C. difficile* has multiple metabolic pathways that overlap to ensure generation of key metabolites. The Wood-Ljungdahl pathway was recently found to be not as active in *C. difficile* as in other acetogens, but this pathway can be used in conjunction with butyrate formation to regenerate NAD^+^ in the absence of Stickland metabolism (13). Carbon metabolism includes breaking down sugars such as glucose and mannitol to generate pyruvate and acetyl-CoA for glycolysis and the tricarboxylic acid cycle, although the latter is incomplete. Pyruvate is a key metabolite in many different central carbon metabolism and fermentation pathways (12). It can be utilized by pyruvate formate-lyase to generate CO_2_ for carbon fixation (14, 15), and it can be degraded to acetyl-CoA to generate butyrate (12). Pyruvate is also used in fermentation pathways to generate propionate via the reductive branch of Stickland metabolism. Fermentation pathways also include Stickland metabolism which contributes to electron bifurcation and the membrane spanning Rnf complex to generate a sodium/proton gradient for substrate-level phosphorylation. While it appears that many of the metabolic processes in *C. difficile* have been elucidated, how these processes interact when others are impaired has yet to be studied (12).

Stickland metabolism is a primary source of energy for a small group of anaerobic bacteria which use amino acids as their sole carbon and nitrogen source (*i.e., C. difficile, Clostridium sporogenes*, and *Clostridium sticklandii*). The main goal of this metabolic pathway is to generate NAD^+^ and a small amount of ATP for the cell (16–19). In the oxidative branch, an amino acid, most frequently isoleucine, leucine or valine (ILV), is decarboxylated or deaminated and generate products for other metabolic processes + NADH (17, 19, 20). In the reductive branch, D-proline or glycine are deaminated or reduced by their respective reductases (proline reductase, PrdB, and glycine reductase, GrdA) to regenerate NAD^+^ to be reused by the cell (17, 18, 21, 22). Recently, Stickland metabolism was suggested as a significant contributor during *C. difficile* infection in a murine model. Proline and hydroxyproline were found to be the most abundant molecules at the start of infection; 5-aminovalerate, a product of the proline reductase in Stickland metabolism, is an abundant molecule towards the end of infection (23, 24).

Both the proline and glycine reductases are selenoproteins (20, 25). Selenoproteins are made through the incorporation of selenium, as selenocysteine, during protein synthesis. Selenocysteine is generated through a synthesis pathway where inorganic phosphate reacts with hydrogen selenide to generate selenophosphate by the selenophosphate synthetase, SelD. Through the use of a selenocysteinyl-tRNA (Sec) synthase, SelA, selenophosphate is incorporated into serine-charged tRNAs to generate selenocysteine. The selenocysteine-specific elongation factor, SelB, recognizes an in-frame stop codon followed by a selenocysteine insertion sequence (SECIS). This recognition results in a halt in translation to allow for the incorporation of the selenocysteine into the protein sequence (26).

We hypothesized that if we eliminated the global production of selenoproteins, the two Stickland reductases would not be generated, and the resulting strain would be incapable of performing Stickland metabolism. Previously, we generated a CRISPR-Cas9 genome editing tool for use in *C. difficile* (27). In that work, we created a *C. difficile* Δ*selD* strain and analyzed the growth phenotype of this strain compared to the wild-type parent and the mutant complemented with a wild-type *selD* allele *in trans*. We showed that *C. difficile* R20291 Δ*selD* (KNM6) had no growth defect in rich BHIS medium but did have a slight growth defect in TY medium, which is a peptide rich medium and should encourage the cells to use Stickland metabolism for growth (27). Here, we build upon our prior work by using the CRISPR-Cas9 system to restore the *selD* gene at its native locus. Using this *selD*-restored strain, we sought to further characterize this mutant throughout different lifecycles to determine the global role of selenoproteins on *C. difficile* physiology. We find that loss of selenophosphate / Stickland metabolism reprograms *C. difficile* metabolism to pathways that can regenerate NAD^+^ from NADH.

## Results

### Complementation *in trans* results in growth differences at different hydrogen levels

When originally characterizing the *C. difficile* KNM6 (Δ*selD*) mutant strain, we found that the complementing plasmid would not restore the mutant to growth comparable to wild-type, in either TY or TYG medium, despite the use of a fragment upstream of the *selDAB* operon that should contain the native promoter region (Figure 1A). However, as published in the prior work, we were able to complement the phenotype, and this complementation was dependent on the abundance of hydrogen gas in the anaerobic chamber. When monitored using a COY Anaerobic Monitor (CAM-12), a 4% hydrogen level did not permit the complementing plasmid to restore the growth phenotype to wild-type levels (Figure 1A). However, when the hydrogen level was lowered to ~1.7% we observed complementation in TY medium (Figure 1B). This observation was interesting and suggests that hydrogen abundance in the anaerobic chamber influences *C. difficile* physiology or the ability of *selD*, when expressed from a multicopy plasmid, to function within *C. difficile*. However, to fully characterize the *C. difficile* Δ*selD* mutant, we wanted to avoid this problem altogether by restoring the CRISPR-Cas9-mediated deletion with *selD* at its native locus.

**Figure 1:**
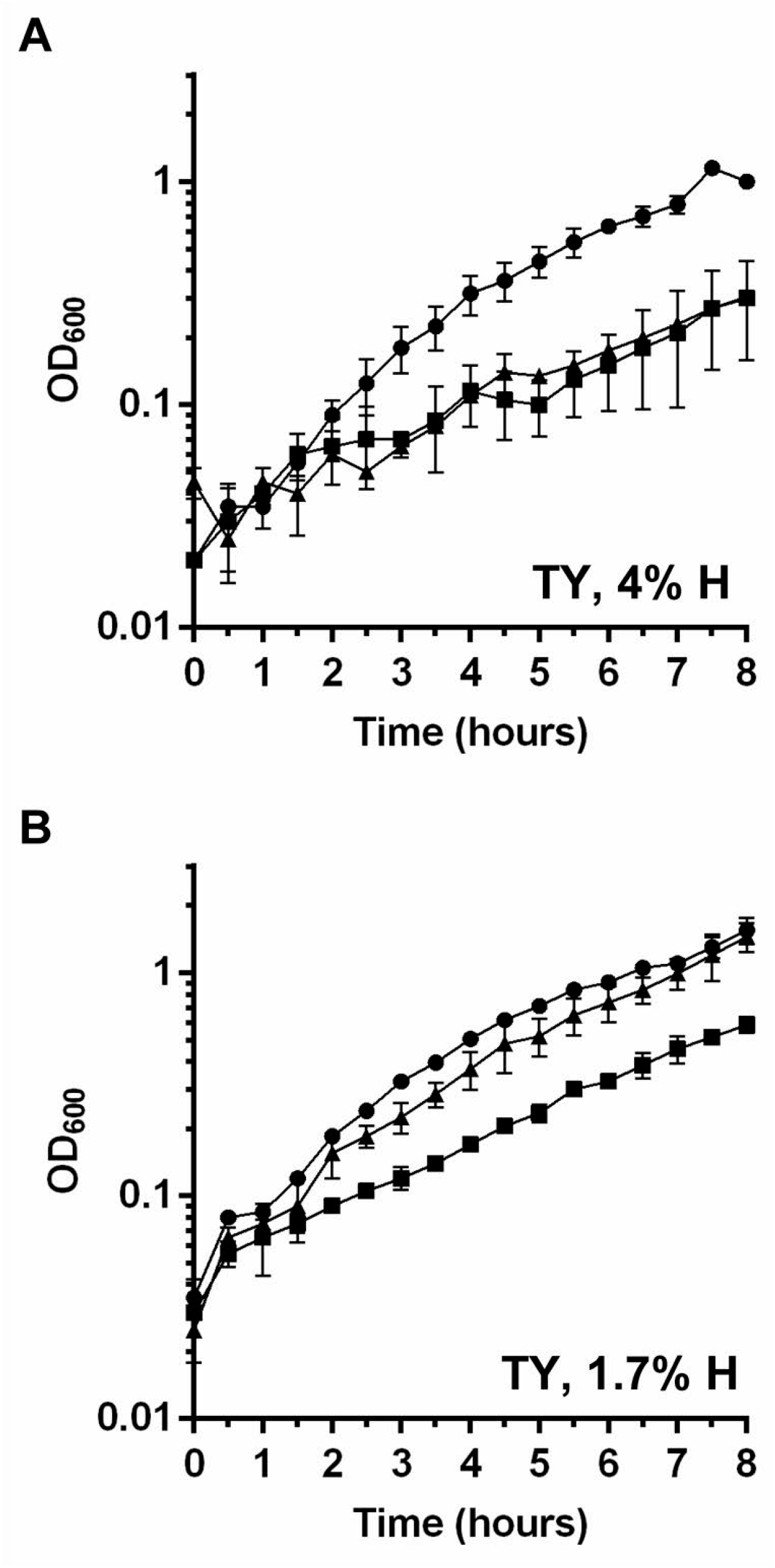
Growth curves with complementing plasmid. *C. difficile* R20291 pJS116 (wild-type, empty vector) (●), *C. difficile* KNM6 pJS116 (Δ*selD*, empty vector) (■), and *C. difficile* KNM6 pKM142 (Δ*selD*::*selD^+^*, p*selD*) (▲) were grown in TY medium at (A) 4% hydrogen and (B) 1.7% hydrogen and growth was monitored over an 8 hour period. Data points represent the average from two independent experiments and error bars represent the standard deviation from the mean.

### Generation of a *C. difficile* restored *selD*^+^ strain by CRISPR-Cas9 genome editing

Our previously developed tool was used to generate a Δ*selD* strain and was limited to this type of mutation. By modifying the existing CRISPR-Cas9 plasmid, we have improved the functionality of this tool to include insertions within the genome. Recently, Muh *et al*. developed a *C. difficile* CRISPRi tool which included dCas9 to be under the conditional expression of a xylose-inducible promoter system (28). We replaced the previous tetracycline-inducible promoter system, which was shown to have uncontrolled expression of Cas9, with the xylose-inducible promoter. We replaced the homology region and gRNA target sequences in our new CRISPR-Cas9 gene editing plasmid. The targeting region is 26 bp away from the deletion site to reintroduce *selD* at its native locus (Figure 2A). Restoration-plasmid-containing *C. difficile* Δ*selD* was passaged on xylose-containing agar medium. Using this strategy, we generated a *C. difficile selD* restored strain (KNM9; Δ*selD*::*selD*^+^) (Figure 2B).

**Figure 2:**
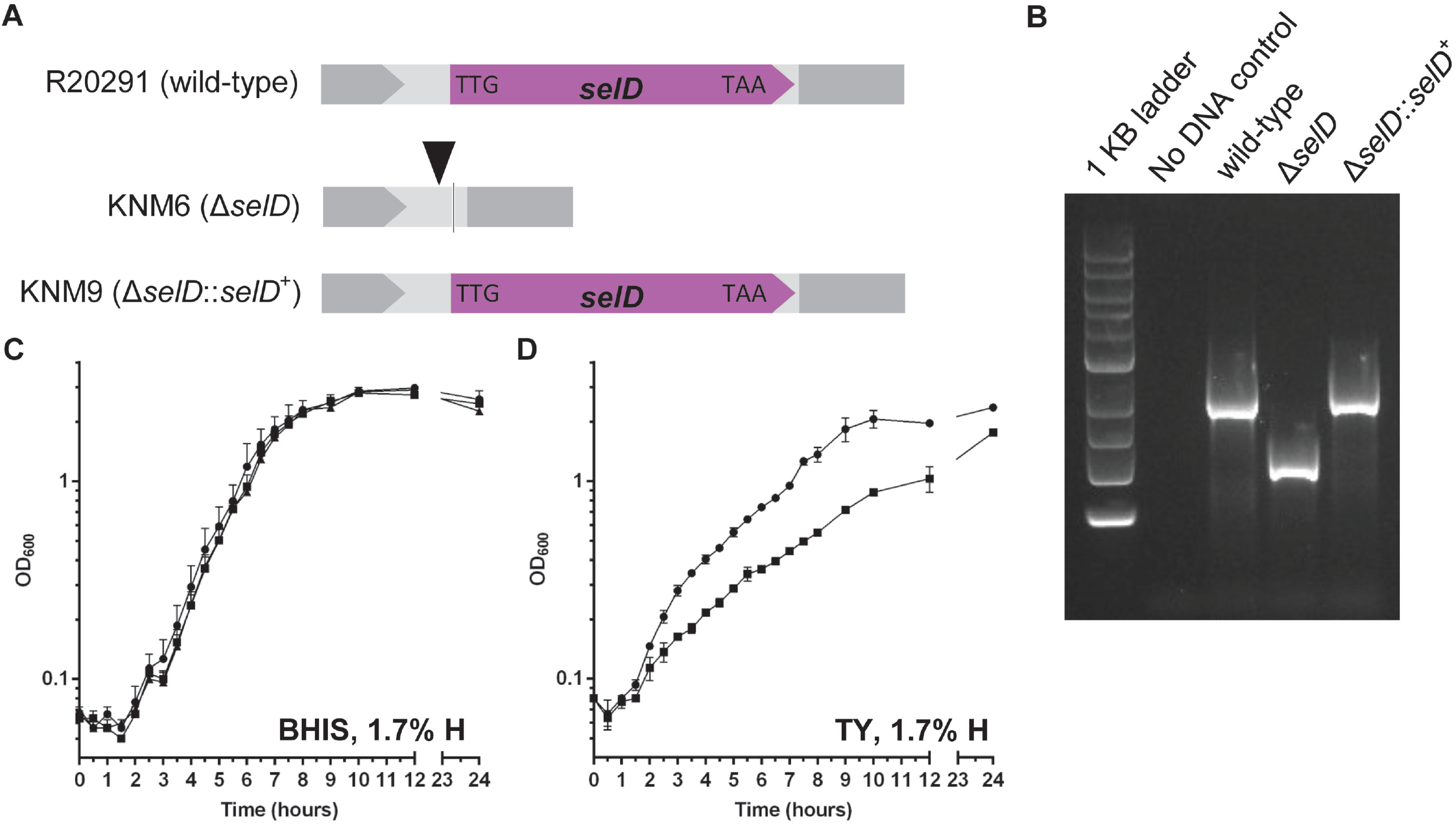
Insertion by *C. difficile* CRISPR-Cas9 genome editing tool and the slight growth defect of a *C. difficile* Δ*selD* strain. (A) Graphical representations of the three strains used in this study. Shown as a triangle is the target site of the gRNA and the line indicates the site of the deletion. (B) DNA was isolated from *C. difficile* R20291 (wild-type), KNM6 (Δ*selD*), and KNM9 ((Δ*selD*::*selD*^+^). The region surrounding the *selD* gene was amplified from the chromosome, and the resulting DNA was separated on an agarose gel. A clean deletion of *selD* is indicated by a faster-migrating DNA band while wild-type and the insertion mutation (restoration) is indicated by a slower-migrating DNA band. (C-D) *C. difficile* R20291 (wild-type) (●), *C. difficile* KNM6 (Δ*selD*) (■), and *C. difficile* KNM9 (Δ*selD*::*selD*^+^) (▲) were grown in (C) BHIS medium and (D) TY medium at 1.7% hydrogen and growth was monitored over a 24 hour period. Data points represent the average from three independent experiments and error bars represent the standard deviation from the mean.

### *C. difficile* Δ*selD* strain has a growth defect in peptide rich medium

Because we generated a *selD* restored strain, we repeated the growth experiments to confirm that the restored strain would complement the phenotype. When the *C. difficile* wild-type (R20291), Δ*selD* (KNM6), and Δ*selD*::*selD*^+^ (KNM9) strains were grown in rich BHIS medium, we again saw no difference between the growth of these three strains over 24 hours (Figure 2C). When these three strains were grown in TY (Figure 2D) or TYG (Figure S1), we observed a growth defect of the Δ*selD* strain when compared to wild-type or Δ*selD*::*selD*^+^ (Figures 2D and S1). Again, there was no observable difference whether glucose was supplemented in the medium or not on the growth of these strains. For this reason, TY medium was used for most of the subsequent experiments. The growth data are consistent with our previous findings along with the additional finding that the Δ*selD* mutant strain grows to similar levels as the wild-type and restored strains in TY medium at 24 hours (Figure 2D). We will note that these and all subsequent experiments were performed at ~1.7% hydrogen and will discuss our rationale for this later in the manuscript.

### Sporulation

To understand how the absence of selenoproteins impacts *C. difficile* spore formation, we determined the sporulation frequency in *C. difficile* R20291, KNM6 (Δ*selD*) and KNM9 (Δ*selD*::*selD*^+^) (29). As a negative control, we generated a deletion of *spo0A* in *C. difficile* R20291 using CRISPR-Cas9 (Figure S2). When compared to the *C. difficile* R20291, wild-type, strain, the *C. difficile* Δ*selD* mutant produced fewer spores in both BHIS and TY media (Figure 3A). Restoration of the *selD* gene resulted in a return of sporulation to wild-type levels. These results indicate that the absence of selenoproteins / selenophosphate has a mild impact on *C. difficile* sporulation.

**Figure 3:**
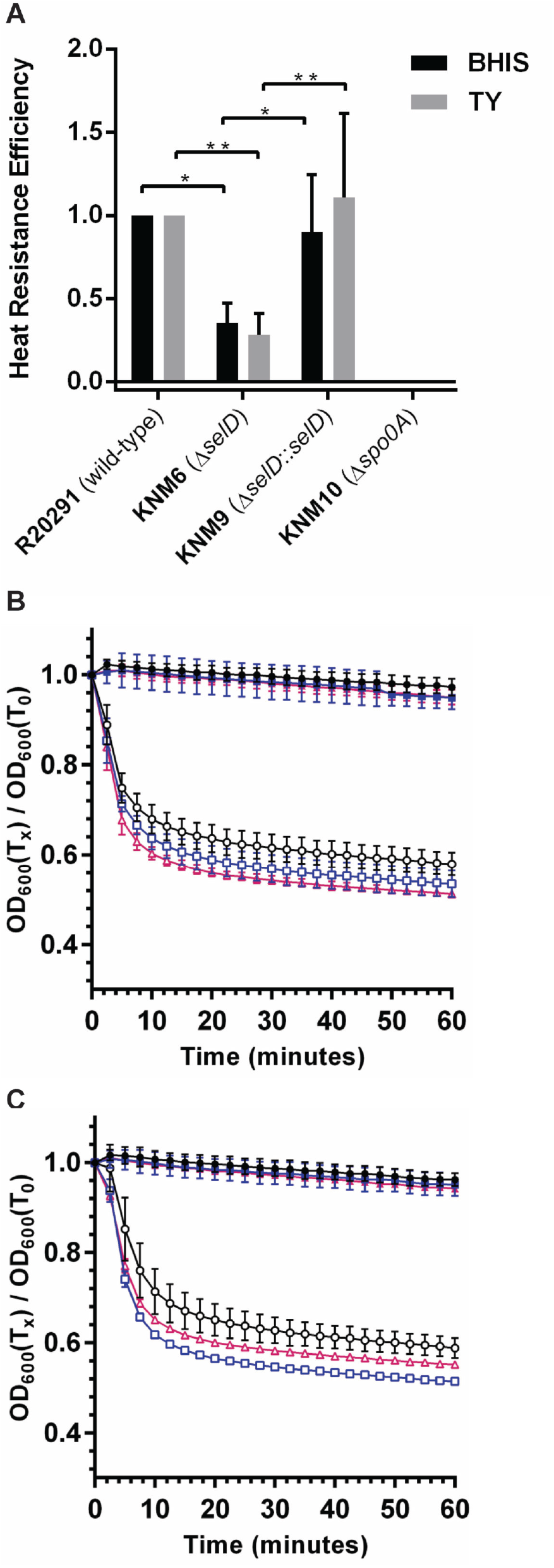
Selenoproteins do not have an effect on germination of *C. difficile* R20291 spores. Purified *C. difficile* R20291 (wild-type) (●/○), *C. difficile* KNM6 (Δ*selD*) 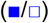, and *C. difficile* KNM9 (Δ*selD*::*selD*^+^) 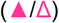 spores were suspended in (A) BHIS or (B) TY medium supplemented with 10 mM taurocholate (open symbols) or without taurocholate (closed symbols). The change in OD_600_ during germination was measured over time at 37°C. The data represents the average of three biological replicates and error bars represent the standard deviation from the mean.

### Selenoprotein synthesis has no effect on *C. difficile* spore germination

Due to the mild defect in sporulation, we then wanted to determine whether the Δ*selD* strain had any defect in germination. Strains were grown and allowed to sporulate on BHIS medium to minimize any affects from both growth and sporulation defects. The spores were purified and analyzed for germination using the optical density assay. When *C. difficile* wild-type (R20291), Δ*selD* (KNM6), and Δ*selD*::*selD*^+^ (KNM9) spores were suspended in rich BHIS medium supplemented with 10 mM taurocholate (TA), rapid germination occurred (Figure 3B). Since the mutant phenotype is apparent in TY medium, we germinated spores in this medium supplemented with 10 mM TA and measured the drop in optical density. Similar to rich medium, the spores rapidly germinated in TY medium (Figure 3C). These results suggest that the selenophosphate synthetase, *selD*, plays no significant role in the early events during spore germination.

### Significant delay in outgrowth of Δ*selD* germinated spores in peptide-rich medium

Because we observed a defect in growth of the *C. difficile* KNM6 (Δ*selD*) strain, we hypothesized that the strain would have a deficiency in outgrowth from spores. Purified spores from wild-type *C. difficile* R20291, KNM6 (Δ*selD*), and KNM9 (Δ*selD*::*selD*^+^) were germinated for 10 minutes at 37°C in rich BHIS medium to ensure that the germination conditions would not impact outgrowth of a vegetative cell from the germinated spore. After washing the germinated spores in either BHIS or TY medium, the germinated spores were resuspended in the anaerobic chamber in pre-reduced BHIS or TY medium. We then monitored the optical densities over a 24 hour period (Figure 4). When cells were allowed to outgrow in rich BHIS medium, there was a non-significant delay in growth in the *C. difficile* KNM6 (Δ*selD*) strain of approximately thirty minutes, when compared to the wild-type and restored strains (Figure 4A). On the other hand, when cells were allowed to outgrow in peptide-rich, TY medium, *C. difficile* KNM6 (Δ*selD*) had an observed outgrowth defect, compared to the wild-type and restored strains (Figure 4B).

**Figure 4:**
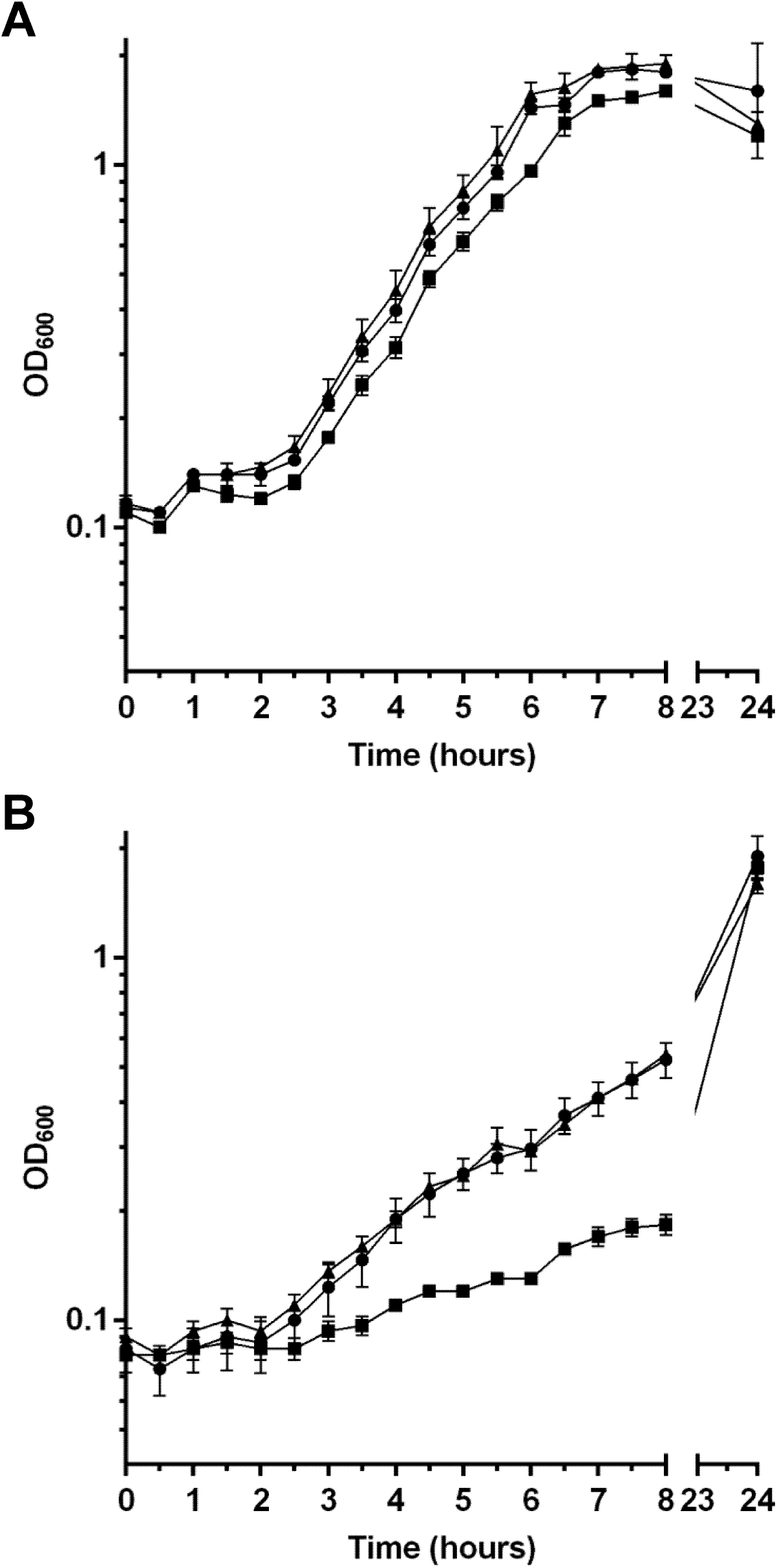
*selD* plays a role in outgrowth of *C. difficile* R20291 spores. Purified *C. difficile* R20291 (wild-type) (●), *C. difficile* KNM6 (Δ*selD*) (■), and *C. difficile* KNM9 (Δ*selD*::*selD*^+^) (▲) spores were germinated and then resuspended in (A) BHIS or (B) TY medium, and the OD_600_ was measured over an 8 hour period and then again at 24 hours. Data points represent the average from three independent experiments and error bars represent the standard deviation from the mean.

### RNA-seq comparison of wild-type versus Δ*selD* strains

If the hypothesis that Stickland metabolism is a primary source of energy for *C. difficile* is true, we wondered why the growth defect in peptide-rich medium was not more severe compared to wild-type and restored strains. We then hypothesized that the *C. difficile* KNM6 (Δ*selD*) strain was able to compensate for the loss of global selenoprotein synthesis through upregulation of other metabolic pathways. To determine which pathways were differentially expressed, we performed RNA-seq on the wild-type *C. difficile* R20291 and *C. difficile* KNM6 (Δ*selD*) strains during exponential growth in TY medium supplemented with glucose. At the time of performing these experiments, we were not aware of whether glucose would have an effect on gene expression and, therefore, decided to include this carbohydrate in the medium. We performed these experiments at both ~1.7% and 4% hydrogen levels to determine whether atmospheric hydrogen abundance impacts *C. difficile* physiology.

First, we noticed when comparing expression of genes in wild-type cells grown at low and high hydrogen levels, the cells had significant down-regulation of ribosomal proteins at high hydrogen. This suggests that the cells perceive high hydrogen as a stressful condition. Due to this finding, we chose to exclude highly up- or down-regulated gene expression due to hydrogen levels when comparing wild-type and mutant expression levels. Meaning, we excluded expression of genes which had significant expression changes in either low or high hydrogen levels of wild-type cultures. After this exclusion, we obtained a more narrow list for comparison of wild-type and Δ*selD* strains at low hydrogen levels and further determined their known or hypothesized function and what pathway that gene belongs (Table 1) (30, 31). As expected, many of the genes are involved in metabolism; 36.35% of genes which were up-regulated, and 22.24% of genes which were down-regulated (Figure 5). In both cases, there were large numbers of genes of unknown function, more than 25% both up- and down-regulated. Other than metabolism and unknown functions, the largest group which contained up-regulated expression in the Δ*selD* strain characterized as transferases. Although, many of these genes also fit into another KEGG pathway, *i.e*. metabolism. Another group of genes which was significant were those of transporters or involved in secretion, likely increasing the cell’s import and export systems to compensate for this growth impairment. The largest group of genes which were down-regulated, besides metabolism or unknown functions, belonged to secretion systems or hydrolases. Again, the cell was likely increasing certain pathways or import / export systems to allow energy to be utilized elsewhere.

**Table 1:**
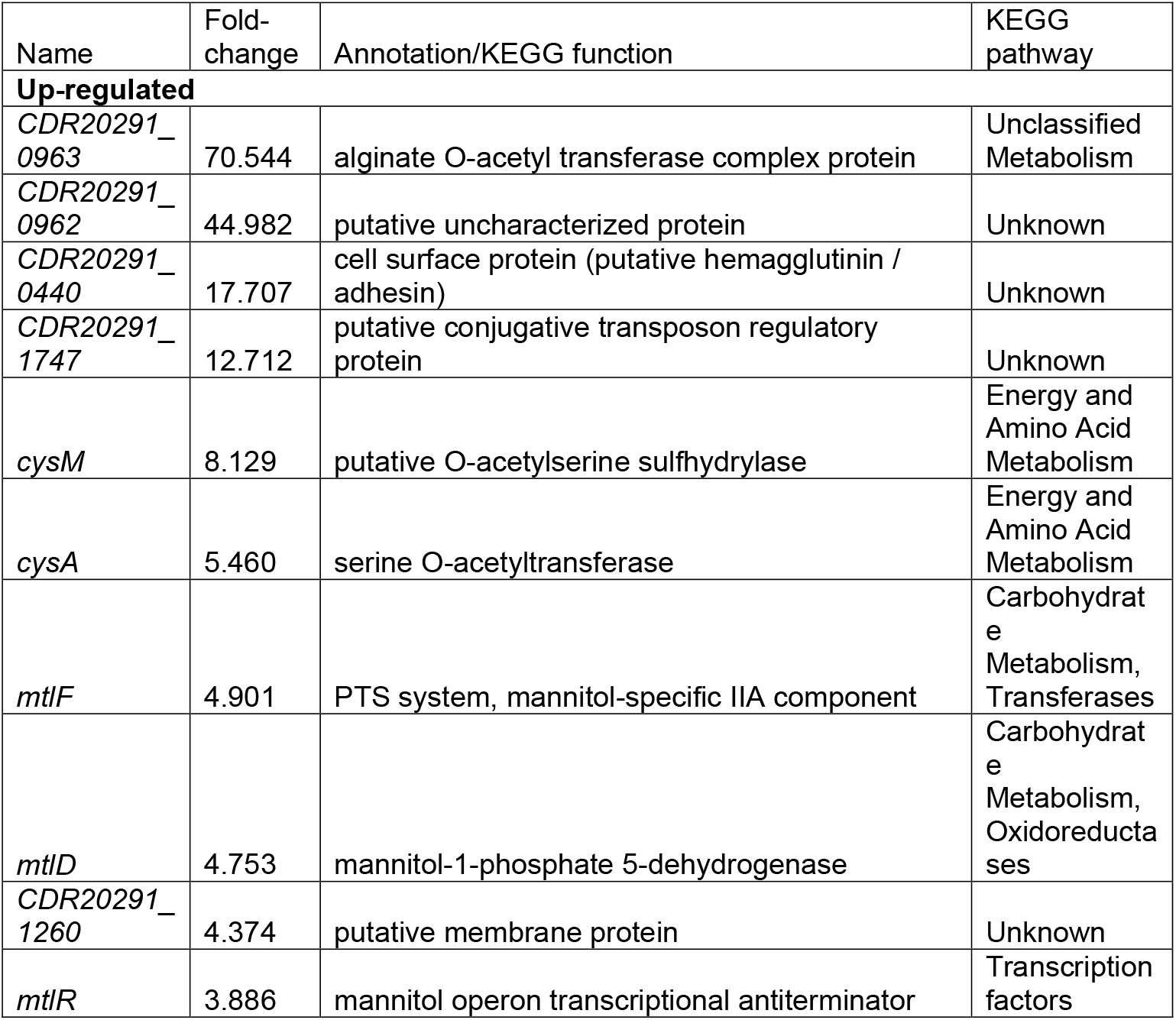
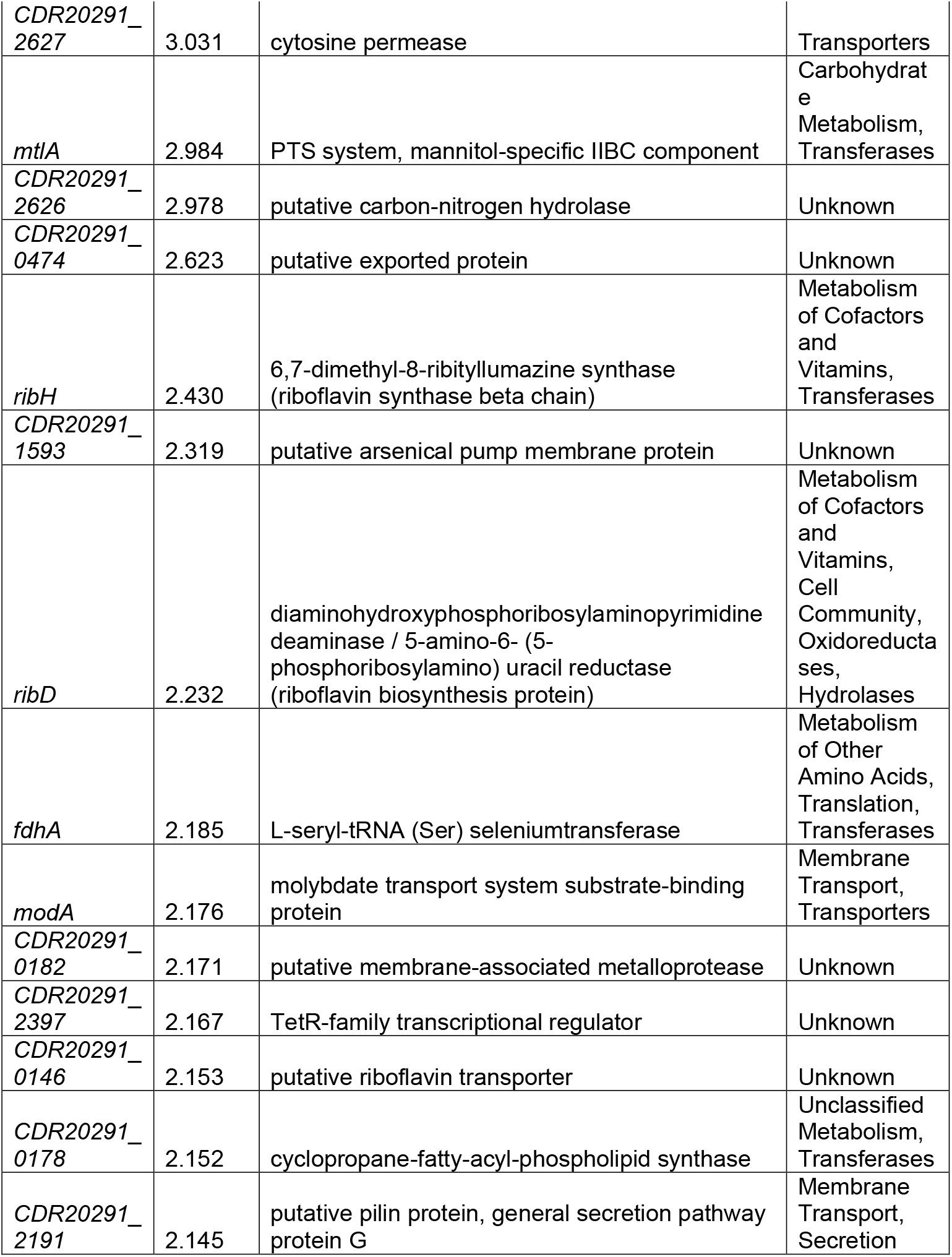
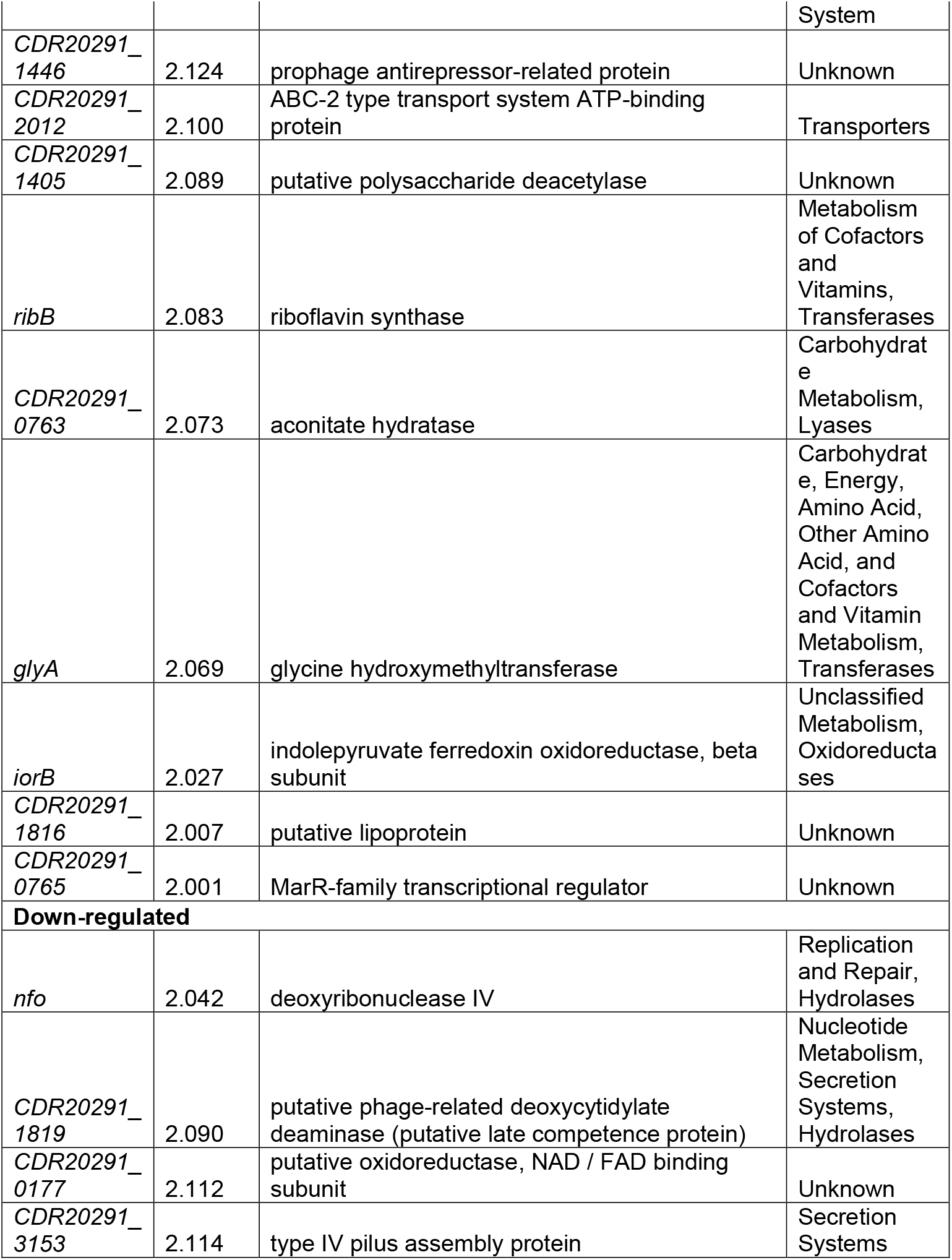
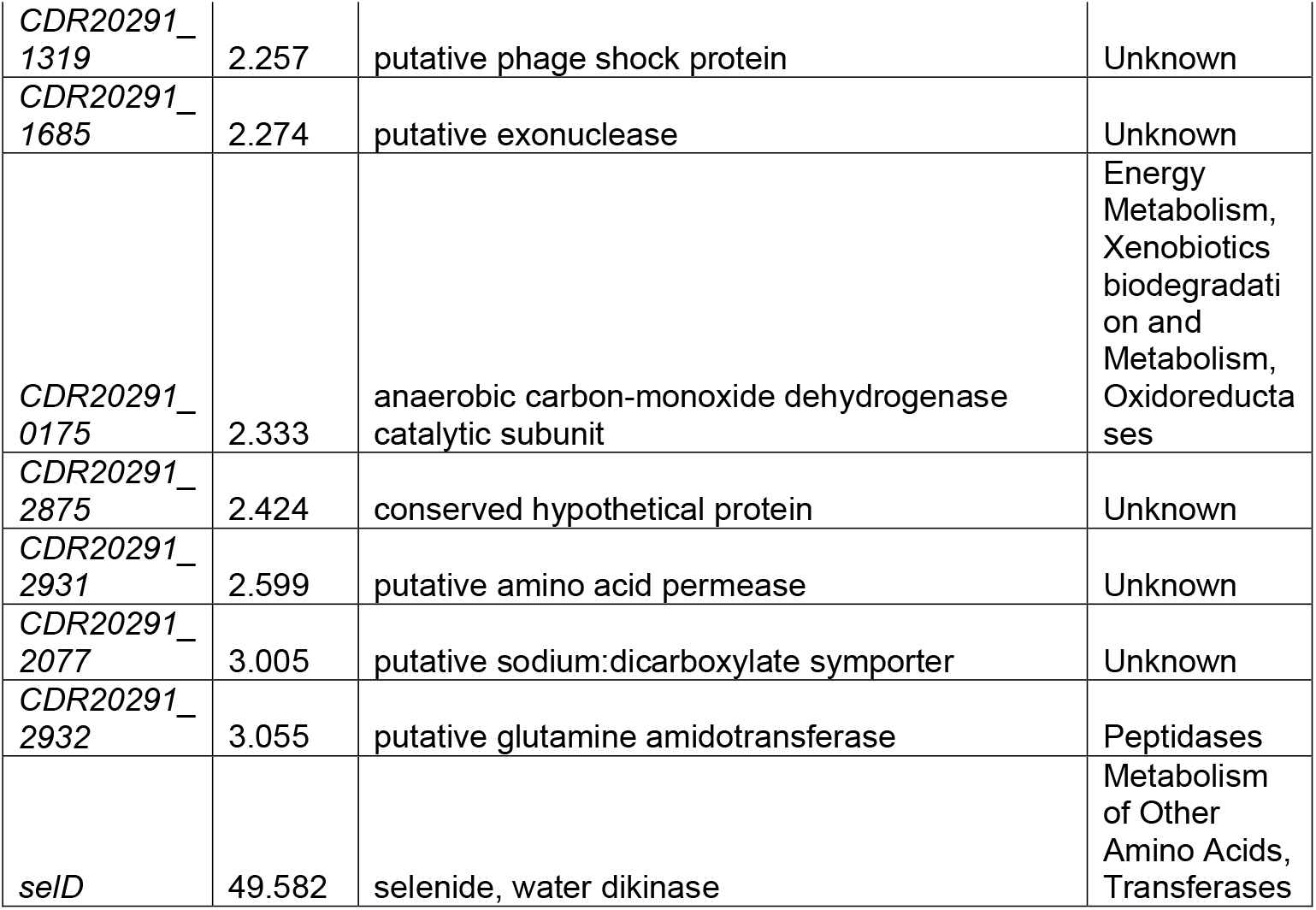
Differentially expressed genes in the *C. difficile* Δ*selD* strain compared to the wild-type strain.

**Figure 5:**
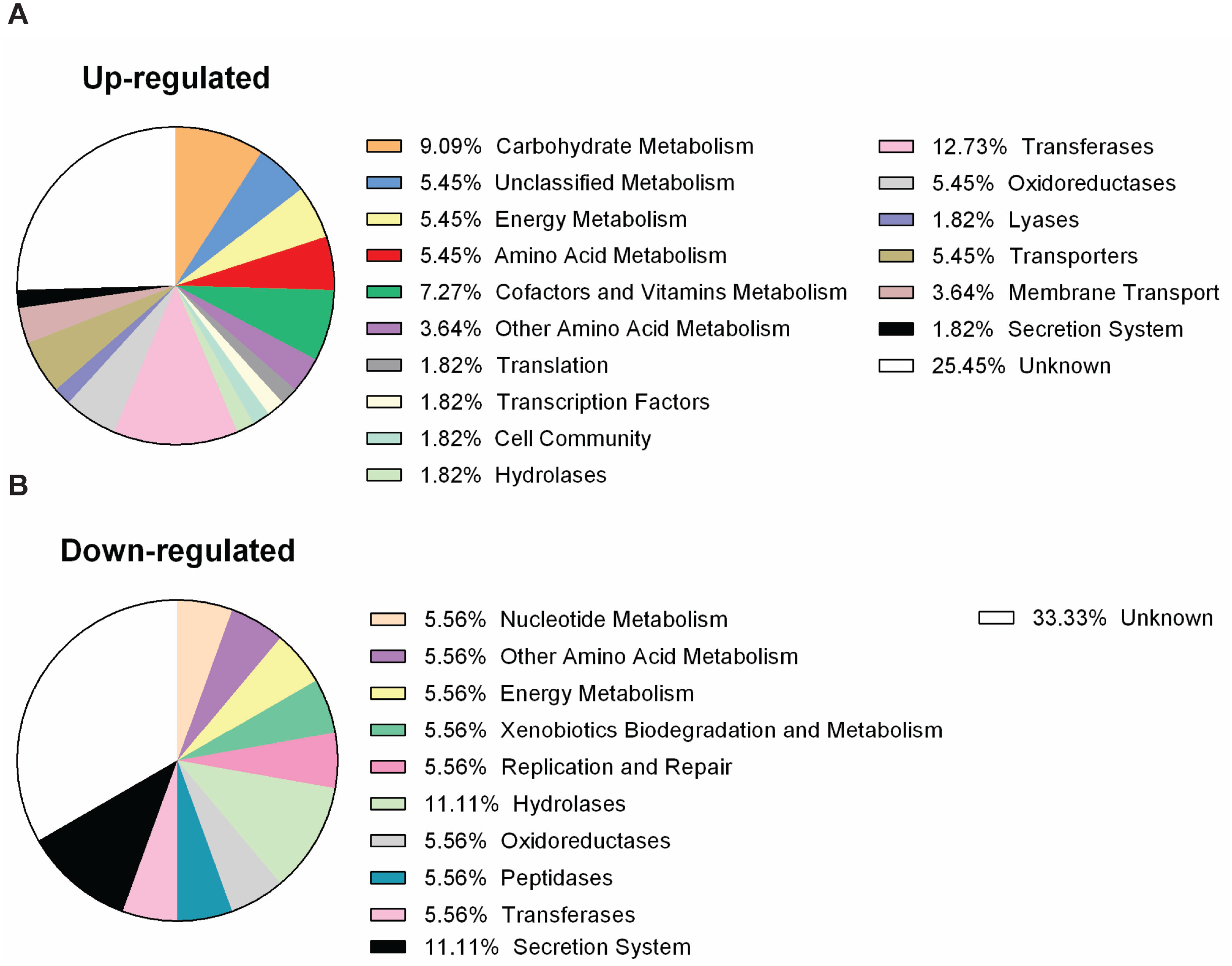
Distribution of functions of up- and down-regulated genes from RNA-seq. Pie charts of the KEGG functional pathways of genes which were (A) up-regulated and (B) down-regulated from RNA-seq analysis. Percentages of each function are shown in the legends.

### Validation of RNA-seq by quantitative RT-PCR

We then chose genes which we have highlighted as well as others of interest to validate the RNA-seq results by qRT-PCR. *C. difficile* R20291 and *C. difficile* KNM6 strains were grown as previously described for RNA-seq, DNA was depleted, and cDNA libraries were generated. Using the housekeeping gene, *rpoB*, as an internal normalization control, we determined the fold change of transcripts in the KNM6 (Δ*selD*) strain compared to the R20291 (wild-type) strain at both 4% and 1.7% hydrogen levels. The target genes *CDR20291_0962* and *CDR20291_0963* had transcript levels which were interesting at 1.7% hydrogen percentage since it appeared that, in one biological replicate, the riboswitch was likely turned on while other biological replicates had fold change values close to one and were likely turned in the off position (Figure 6A and 6B). One target gene, *mtlF*, which is part of the mannitol utilization pathway, had a slightly increased fold change in transcript levels at both 4% and 1.7% by qRT-PCR, and the trend correlated to the results found from RNA-seq (Figure 6C). Mannitol utilization may be a large factor helping the strain grow in absence of selenoproteins.

**Figure 6:**
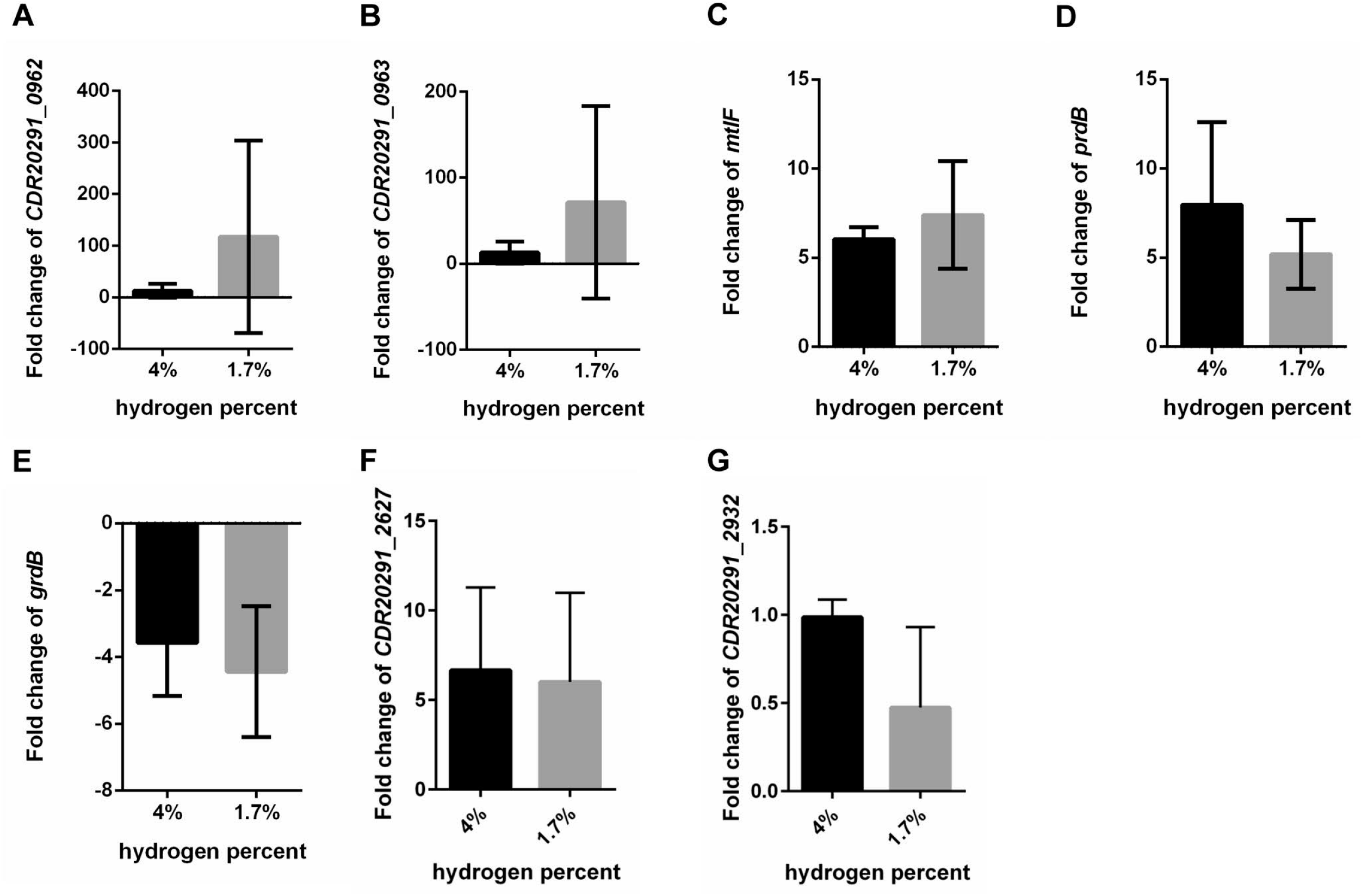
Fold change of transcript levels in *C. difficile* KNM6 (Δ*selD*) compared to *C. difficile* R20291. RNA was extracted from *C. difficile* R20291 and KNM6 (Δ*selD*) strains grown to an OD_600_ of 0.6 in TYG medium, DNA was depleted, and cDNA was synthesized. Quantitative reverse transcription-PCR was performed to determine the fold change of transcript levels in the KNM6 (Δ*selD*) strain compared to wild-type R20291 strain at each 4% and 1.7% hydrogen levels. Fold change for the following genes are shown: (A) *CDR20291_0962*, (B) *CDR20291_0963*, (C) *mtlF*, (D) *prdB*, (E) *grdB*, (F) *CDR20291_2627*, and (G) *CDR20291_2932*. Data represents the average from three independent experiments run in technical triplicate and error bars represent the standard deviation from the mean. Technical replicate outliers were excluded.

Since we hypothesized Stickland metabolism to be impaired in the mutant strain, we chose to analyze the proline reductase, *prdB*, and the glycine reductases, *grdB*. The fold change in expression of *prdB* was higher than was seen in RNA-seq (Figure 6D). In qRT-PCR, the transcription of *grdB* was down-regulated in the mutant compared to wild-type (3.6- and 4.4-fold down-regulated in 4% and 1.7% hydrogen, respectively) (Figure 6E). On the other hand, the transcription of *grdB* was slightly up-regulated in RNA-seq (1.3- and 2-fold up-regulated in 4% and 1.7% hydrogen, respectively).

Finally, we validated two uncharacterized genes. *CDR20291_2627*, a putative hemagglutinin / adhesion, was up regulated in the RNAseq at both 4% hydrogen and 1.7% hydrogen. We also observed upregulation of this gene in the qRT-PCR validations. Additionally, *CDR20291_2932*, a putative glutamine deaminase, was slightly downregulated, and we validated this by qRT-PCR.

## Discussion

It is becoming increasingly appreciated that Stickland metabolism and Stickland substrates are important for *C. difficile* pathogenesis. In the oxidative branch of Stickland metabolism, several different amino acids can be deaminated or decarboxylated. In the reductive branch either proline or glycine are used by their respective reductases. The proline reductase, PrdB, converts proline + NADH to 5-aminovalerate + NAD^+^ while the glycine reductase, GrdA, converts glycine + NADH + ADP to acetate + ATP + NAD^+^. The activity of the PrdB and GrdA proteins is dependent on the incorporation of a modified amino acid, selenocysteine.

Incorporation of selenium into proteins is governed by the SelD, SelA and SelB proteins. The first step in this pathway is the generation of selenophosphate from hydrogen selenide by the selenophosphate synthetase, SelD. We hypothesized that a mutation in *C. difficile selD* would significantly alter *C. difficile* physiology by blocking the total incorporation of selenium into protein as an amino acid, selenocysteine, or as part of selenium containing cofactors. Surprisingly, we found that plasmid-based complementation of the *selD* deletion led to an odd observation – that complementation only occurred at low hydrogen percentage (1.7%) in our anaerobic chamber. At higher hydrogen abundance (4%), we could not complement our mutant.

To address this, we used our CRISPR/Cas9 genome editing tool to restore the *selD* gene at its native locus. By targeting the Cas9 nuclease to a region just outside of the deleted region, we were able to isolate a restored strain. This *C. difficile* Δ*selD*::*selD*^+^ strain grew similar to the wild-type strain. We then analyzed the physiology of the strains. We observed no differences in germination between the wild-type, mutant and restored strains, but the Δ*selD* strain had a mild decrease in spore formation. However, we observed a significant delay during the outgrowth of germinated *C. difficile* spores in peptide-rich medium (TY). This highlights the importance of selenophosphate during the early stages of return to vegetative growth.

To understand the overall impact of SelD on *C. difficile* physiology, we determined the global expression differences between the wild-type and mutant strains at low and high hydrogen percentages. First, we observed that *fdhA* was upregulated in the *selD* mutant strain. *fdhA* encodes the selenocysteinyl-tRNA synthetase that charges serine charged tRNA with selenophosphate to generate selenocysteine-charged tRNA. This suggests that in the absence of selenoprotein, *C. difficile* senses this loss and attempts to increase the rate of selenocysteine-charged tRNA.

On first glance, *CDR20291_0963* and *CDR20291_0962* are the two most highly up-regulated genes in the Δ*selD* strain. *CDR20291_0963* is uncharacterized *in C. difficile* but has high similarity to the AlgI protein, an alginate O-acetyltransferase, characterized in *Pseudomonas aeruginosa*. Whether this protein has the same function in *C. difficile* is yet to be determined. The other gene *CDR20291_0962* is an uncharacterized gene of no known function. When ran through NCBI BLAST and the Conserved Domain Database (32, 33), this protein sequence has a DHHW domain which is common among bacteria. What we found interesting is that both of these genes have recently been characterized to contain a riboswitch upstream, and their altered expression is likely unrelated to the *selD* mutation but, rather, due to the riboswitch being in the “ON” position (34).

Interestingly, every gene in the operon *mtlARFD* had upregulated expression in the Δ*selD* strain. The expression levels varied between almost 3- to 5-fold up-regulation compared to wild-type. We hypothesize that mannose metabolism may play a larger role in a strain lacking selenoproteins. In this pathway, mannitol is taken into the cell and broken down to mannitol-1-phosphate by MtlF and MtlA. Then, MtlD uses NAD^+^ to convert mannitol-1-phosphate into fructose-6-phosphate + NADH to then be used for glycolysis (35–38). Prior work by Teschner and Garel (1990) has shown that the *E. coli* MtlD can run this reaction in reverse to generate NAD^+^ instead of consuming it (39).

Besides mannose metabolism, cysteine and methionine metabolism appeared to be increased by the up-regulation of *cysM* and *cysA*. It is interesting that these genes are utilizing H_2_S produced by *C. difficile* to generate L-cysteine and acetate (40, 41). Acetate is also a final product in the reduction of glycine by Stickland metabolism (17). It is possible that the cell is generating acetate through this method in order to utilize the molecule in other metabolic pathways.

Another metabolic process which had multiple genes up-regulated was for riboflavin synthesis: *ribD*, a deaminase and reductase; *ribH*, a lumazine synthase; and *ribB*, a riboflavin synthase. Riboflavin is an essential cofactor which can be broken down into flavins (flavin mononucleotide, FMNs) and flavin adenine dinucleotide (FAD) to be used by the cell. Riboflavin can also be taken up by the environment if present (42). A gene proposed to be a riboflavin transporter, *CDR20291_0146*, was also up-regulated from our RNA-seq analysis. This essential cofactor is likely being used in multiple processes and its synthesis or uptake is being up-regulated to feed these metabolic pathways.

Of those genes which had down-regulated expression, there are a few worth noting. Multiple genes involved in transport or utilization of amino acids were down-regulated, possible due to the decrease in the dependence on Stickland metabolism to provide NAD+ for the cell to generate energy. *CDR20291_2931*, a putative amino acid permease for transport of amino acids into the cell, and *CDR20291_2932*, a putative glutamine amidotransferase for transforming glutamine into generating an available carbon-nitrogen group, both are proposed to be involved in the utilization of amino acids (43, 44). As well, a putative sodium:dicarboxylate symporter, *CDR20291_2077*, could also be involved in amino acid uptake into the cell. The down-regulation of these three genes indicates that the cell is moving its resources away from the degradation of amino acids as a source of energy.

Overall, our data highlight the importance of selenophosphate for *C. difficile* physiology. The absence of selenophosphate leads to the rerouting of *C. difficile* metabolism so that the cell is less dependent on Stickland metabolism of amino acids and the regeneration of NAD^+^ by the reductive branch of Stickland. However, what was surprising was the impact of atmospheric chamber conditions (H_2_ percentage) on *C. difficile* physiology. Further work should be done to understand how hydrogen abundance influences *C. difficile* physiology, potentially in autotrophic metabolism, and when reporting on *C. difficile* physiology the concentration of hydrogen should be reported.

## Materials and Methods

### Bacterial strains and growth conditions

*C. difficile* strains were routinely grown in an anaerobic atmosphere (1.7% - 4% H_2_, 5% CO_2_, 85% N_2_) at 37°C in brain heart infusion supplemented with 5 g / L yeast extract and 0.1% L-cysteine (BHIS), as described previously (8, 45–47), or TY media (3% tryptone, 2% yeast extract) (48). Hydrogen levels were determined by a COY Anaerobic Monitor (CAM-12). For conjugation experiments, cells were plated on TY medium for *Bacillus subtilis-based* conjugations. Where indicated, growth was supplemented with taurocholate (TA; 0.1% w / v), thiamphenicol (10 μg / mL), D-cycloserine (250 μg / mL), xylose (1% w / v) and / or glucose (1% w / v) as needed. Induction of the CRISPR-Cas9 system was performed on TY agar plates supplemented with thiamphenicol (10 μg / mL) and xylose (1% w / v). *E. coli* strains were routinely grown at 37°C in LB medium. Strains were supplemented with chloramphenicol (20 μg / mL) as needed. *B. subtilis* BS49 was routinely grown at 37°C in LB broth or on LB agar plates. Strains were supplemented with chloramphenicol (2.5 μg / mL) and/or tetracycline (5 μg / mL).

### Plasmid construction and molecular biology

To construct the CRISPR-Cas9 *selD* complementing plasmid, the previously published CRISPR-Cas9 *pyrE* targeting plasmid,pJK02 (27), was modified by replacing *traJ* with oriT *tn916* for *B. subtilis* conjugation by amplification from pJS116 using primers 5’Tn916ori and 3’Tn916ori. The resulting fragments was introduced into pJK02 by Gibson assembly at the *Apa*I site and transformed into *E. coli* DH5α to generate pKM126. Next, the donor region to be used for homology directed repair was PCR amplified from *C. difficile* R20291 genomic DNA using primers 5’selD_comp and 3’selD_comp 2 where this fragment contains a 500 bp upstream homology arm and a 500 bp downstream homology arm surrounding *selD*. The resulting fragment was cloned by Gibson assembly into pKM126 at the *Not*I and *Xho*I restriction sites and transformed into *E. coli* DH5α to generate pKM181. Lastly, the gBlock for *selD* targeting sgRNA, CRISPR_selD_comp2, was introduced by Gibson assembly into the *Kpn*I and *Mlu*I restriction sites and transformed into *E. coli* DH5α resulting in pKM183. To improve efficiency of the CRISPR-Cas9 editing system and provide more control of Cas9 expression, the tetracycline-inducible system was replaced with the xylose inducible system (28). To do this, the xylose inducible promoter was PCR amplified from pIA33 using primers 5’selDcomp_HR_xylR 2 and 3’cas9_Pxyl 2 and inserted by Gibson assembly into pKM183 at the *Xho*I and *Pac*I restriction sites and transformed into *E. coli* DH5α to generate pKM194.

To construct the CRISPR-Cas9 *spo0A* deletion mutant, the gBlock for *spo0A* targeting sgRNA, CRISPR_spo0A_2, was introduced by Gibson assembly into the *Kpn*I and *Mlu*I restriction sites in pKM197 (49) and transformed into *E. coli* DH5α resulting in pKM213. The homology arms to be used for homology directed repair was PCR amplified from *C. difficile* R20291 genomic DNA using primers 5’spo0A_UP and 3’spo0A_UP for the 500 bp upstream arm and primers 5’spo0A_DN and 3’spo0A_DN for the 500 bp downstream arm. The resulting fragments were then cloned by Gibson assembly at the *Not*I and *Xho*I restriction sites and transformed into *E. coli* DH5α resulting in pKM215.

### Conjugation for CRISPR-Cas9 plasmid insertion

CRISPR-Cas9 plasmid pKM194 was transformed into *B. subtilis* BS49 to be used as a donor for conjugation with *C. difficile* KNM6. Likewise, the pKM215 plasmid was transformed into *B. subtilis* BS49 to be used as a donor for conjugation with *C. difficile* R20291. *C. difficile* R20291 or KNM6 was grown anaerobically in BHIS broth overnight. This was then diluted in fresh pre-reduced BHIS broth and grown anaerobically for 4 hours. Meanwhile, *B. subtilis* BS49 containing the CRISPR plasmids were grown aerobically at 37°C in LB broth supplemented with tetracycline and chloramphenicol for 3 hours. One hundred microliters of each culture was pated on TY agar medium. After 24 hours, the growth was harvested by suspending in 2 mL pre-reduced BHIS broth. A loopful of this suspended growth was spread onto several BHIS agar plates supplemented with thiamphenicol, kanamycin, and D-cycloserine. *C. difficile* transconjugants were screened for the presence of Tn916 using tetracycline resistance, as described previously. Thiamphenicol-resistant, tetracycline-sensitive transconjugants were selected and used for further experiments.

### Induction of the CRISPR-Cas9 system and isolating mutants

The *C. difficile* KNM6 strain containing the *selD*-targeting plasmid, pKM194, was streaked onto TY agar medium supplemented with thiamphenicol and xylose for induction. This was then passaged a second time on the same medium. After isolating colonies on BHIS supplemented with xylose, DNA was extracted and tested for the insertion by PCR amplification of the *selD* region using primers 5’selD and 3’selD. From this, one mixed colony out of 12 samples was isolated. This mixed colony was then passaged on TY agar medium supplemented with thiamphenicol and xylose. Colonies were isolated on BHIS agar medium supplemented with xylose, DNA was extracted, and isolates were tested by PCR amplification again as above. From this, 14 colonies were insertions out of 15. Confirmed restored strains were passaged 3 times in BHIS liquid medium in order to lose the CRISPR-Cas9 plasmid. After pick-and-patch on BHIS agar with and without thiamphenicol, loss of plasmid was confirmed by PCR amplification of a portion of *cas9* using primer set 5’tetR_CO_Cas9 and 3’COcas9 (975) and the gRNA using primer set 5’gdh and 3’gRNA 2.

The *C. difficile* R20291 strain containing the *spo0A*-targeting plasmid, pKM215, was streaked onto TY agar medium supplemented with thiamphenicol and xylose for induction. This was then passaged a second time on the same medium. After isolating colonies on BHIS supplemented with xylose, DNA was extracted and tested for the insertion by PCR amplification of the *spo0A* region using primers 5’spo0A_del and 3’spo0A_del. All tested colonies (36 total) were mutants. Two isolates were passaged twice in BHIS liquid medium in order to lose the CRISPR-Cas9 plasmid. After pick-and-patch on BHIS agar with and without thiamphenicol, loss of plasmid was confirmed by PCR amplification of a portion of *cas9* using primer set 5’tetR_CO_Cas9 and 3’COcas9 (975) and the gRNA using primer set 5’gdh and 3’gRNA 2.

### Sporulation and heat resistance assay

To determine differences in sporulation efficiencies between the *C. difficile* R20291 (wild-type), KNM6 (Δ*selD*), KNM9 (Δ*selD*::*selD*^+^) and KNM10 (Δ*spo0A*) strains, sporulation and heat resistance was determined as described previously (29). Briefly, *C. difficile* R20291, KNM6, KNM9, and KNM10 strains were spread onto BHIS agar medium supplemented with taurocholate. From this, colonies were restreaked onto either BHIS or TY agar media, making a lawn on the plate. After 48 hours of growth, half of the plate was harvested and mixed into 600 μL of pre-reduced PBS. Then, 300 μL of the sample was transferred to a separate tube and heat treated at 65°C in a heat block for 30 minutes, inverting the tube every 10 minutes to ensure even heating. Both the untreated and the heat-treated samples were serially diluted in PBS and plated onto BHIS agar medium supplemented with taurocholate. CFUs were counted 22 hours after plating. Heat resistance was calculated by dividing the CFUs for the heat-treated sample by the CFUs for the untreated sample, and the average was calculated for each strain.

### Spore purification

Spores were purified from *C. difficile* R20291, KNM6, and KNM9 strains as previously described (46, 50, 51). Briefly, spores were streaked onto BHIS agar medium (20 – 30 plates) and allowed to sporulate for 5 to 6 days before scraping each into microcentrifuge tubes containing 1 mL of sterile dH_2_O and kept at 4 °C overnight. The spores and debris mixture was washed five times in sterile dH_2_O by centrifuging for 1 minute at 14,000 x g per wash. Spores were combined into 2 mL aliquots in sterile dH_2_O and layered on top of 8 mL of 50% sucrose and centrifuged at 4,000 x g for 20 minutes. The supernatant containing vegetative cells and cell debris was discarded and the spores were resuspended in sterile dH_2_O and washed five more times as before. The spores were stored at 4 °C until use.

### Germination assay

Purified *C. difficile* R20291, KNM6, and KNM9 spores were first heat activated at 65°C for 30 minutes and then placed on ice until use. To compare germination between the three strains, the OD_600_ was measured over time in different media. Germination was carried out in clear Falcon 96-well plates at 37 °C in a final volume of 100 μL and final concentrations of 10 mM taurocholate and 1X BHIS or TY. Spores were added to a final OD_600_ of 0.5 and germination was analyzed for 1 hour using a plate reader (Spectramax M3 Plate Reader, Molecular Devices, Sunnyvale, CA) (50, 52).

### Outgrowth assays

Purified *C. difficile* R20291, KNM6, and KNM9 spores were heat activated at 65°C for 30 minutes and then placed on ice until use. Spore samples were washed one time with dH_2_O and then added spores to a final optical density (OD_600_) of 0.5 in 8 mL of BHIS medium supplemented with taurocholate (10 mM final concentration). Incubated the spores in germination solution in a hot water bath at 37 °C for 10 minutes and then immediately placed on ice. In all subsequent steps, the germinated spores were kept on ice or at 4°C. Washed once with BHIS or TY medium and removed supernatant. The germinated spores were passed into the anaerobic chamber without using vacuum (flooded chamber with gas) and resuspended each in 20 mL of pre-reduced BHIS or TY medium. The outgrowth was monitored by measuring OD_600_ over time.

### RNA processing

*C. difficile* R20291 (wild-type) and KNM6 (Δ*selD*) was grown to an OD_600_ of 0.6 in TYG medium at low (1.7%) and high (4%) hydrogen levels. At that time, RNA was extracted as described using the FastRNA Blue Kit (MP Biologicals). DNA was depleted using the TURBO DNA-free kit (Invitrogen), repeating the steps in the kit three times to achieve complete depletion. rRNA was then depleted using the Ribo-Zero rRNA Removal kit for bacteria (Illumina). Enrichment of mRNA and generation of cDNA libraries were completed using the TruSeq Stranded mRNA library prep kit (Illumina).

### RNA-seq

cDNA libraries were submitted for Illumina high output single-end 50 sequencing at the Tufts Genomic Core. Reads were assembled using the DNASTAR SeqMan NGen 15 program in a combined assembly noting replicates. Raw expression data between wild-type and mutant strains was then normalized to *rpoB* and then quantified Using DNASTAR ArrayStar 15, assemblies were normalized by assigned Reads Per Kilobase of template per Million mapped reads (RPKM), experiments’ values were capped at a minimum of 1, and all genes were normalized by calibration to the median expression value of *rpoB*. Fold-change was determined four ways: wild-type at 4% hydrogen to wild-type at 1.7% hydrogen, wild-type at 1.7% to wild-type at 4%, wild-type at 4% to mutant at 4%, and wild-type at 1.7% to mutant at 1.7%. Expression which was > 2-fold up- or down-regulated from comparison of wild-type at the different hydrogen levels was excluded when comparing the respective wild-type to mutant comparisons. From this, any gene expression which was > 2-fold up- or down-regulated was considered significant. As a final step, we excluded any genes which had little coverage or low reads visualized using DNASTAR GenVision.

### Quantitative RT-PCR

*C. difficile* R20291 (wild-type) and KNM6 (Δ*selD*) was grown, RNA was extracted, and DNA was depleted the same as described above. 50 ng of total RNA was used in cDNA synthesis using the SuperScript III First-Strand Synthesis System (Thermo Scientific) according to the protocol, including controls for each sample without the presence of reverse transcriptase. Primer to be used for quantitative reverse transcription-PCR (qRT-PCR) were designed using the Primer Express 3.0 software (Applied Biosystems) and efficiencies were validated prior to use. cDNA samples were then used as templates for qPCR reactions to amplify *CDR20291_0963, CDR20291_0962, prdB, grdB, mtlF, CDR20291_2627*, and *CDR20291_2932* (primers are listed in Supplemental Table 3) using PowerUp SYBR Green Master Mix (Applied Biosystems) and a QuantStudio 6 Flex Real-Time PCR machine (Applied Biosystems). Reactions were performed in a final volume of 10 μL including 1 μL undiluted cDNA sample and 500 nM of each primer. Reactions were ran in technical triplicate of each biological triplicate for both wild-type and mutant samples at each hydrogen level. Outliers of technical replicate samples were omitted from analysis. Results were calculated using the comparative cycle threshold method (53), in which the amount of target mRNA was normalized to that of an internal control (*rpoB*).

### Statistical Analysis

*C. difficile* R20291 pJS116 (wild-type, empty vector), *C. difficile* KNM6 pJS116 (Δ*selD*, empty vector), and *C. difficile* KNM9 (Δ*selD*::*selD^+^*, p*selD*) strains were grown in biological duplicate. *C. difficile* R20291, KNM6, and KNM9 strains were grown in biological triplicate. *C. difficile* R20291, KNM6, KNM9, and KNM10 strains were allowed to sporulate in biological triplicate. *C. difficile* R20291, KNM6, and KNM9 spores were purified in biological triplicate where each replicate was grown / sporulated and purified separately to be used for germination and outgrowth assays. RNA was extracted from *C. difficile* R20291 and KNM6 strains for RNA-seq in biological duplicate for each strain. RNA was extracted from *C. difficile* R20291 and KNM6 strains for qRT-PCR in biological triplicate and each was run in technical triplicate as well. In each experiment, data represents averages of each of the indicated replicates and error bars represent the standard deviation from the mean. Technical replicate outliers were excluded from qRT-PCR analysis.

## Acknowledgments

We would like to thank Dr. Craig Ellermeier at the University of Iowa for the generous gift of the xylose inducible promoter. We would also like to thank members of the Sorg lab, Dr. Leif Smith, and members from Dr. Leif Smith’s lab at Texas A&M University for their helpful comments and suggestions during the preparation of this manuscript.

This project was supported by awards 5R01AI116895 and 1U01AI124290 to J.A.S. from the National Institute of Allergy and Infectious Diseases. The content is solely the responsibility of the authors and does not necessarily represent the official views of the NIAID. The funders had no role in study design, data collection and interpretation, or the decision to submit the work for publication.

## Supplemental Information Legend

**Figure S1:**
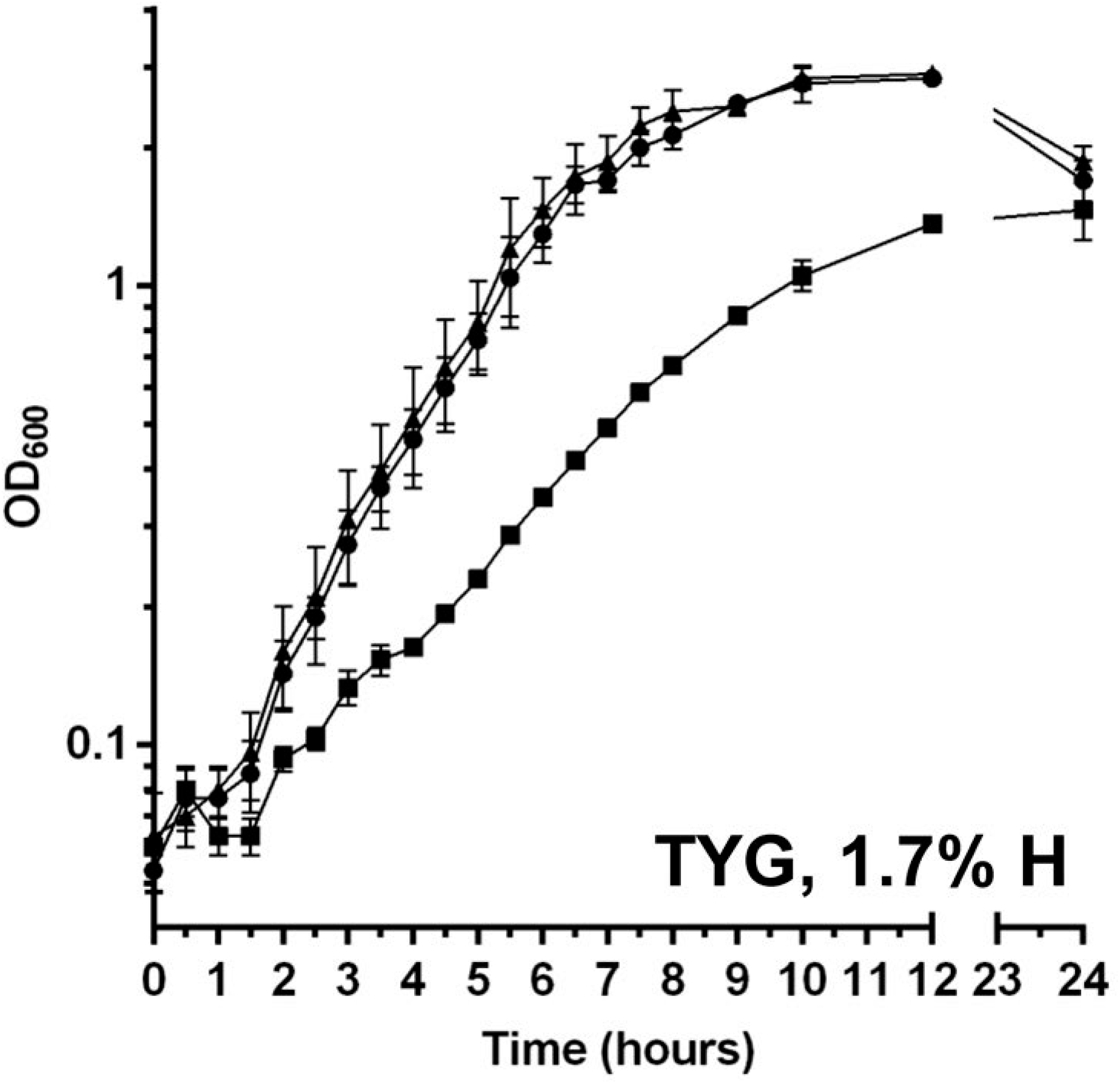
Growth curve in TYG

**Figure S2:**
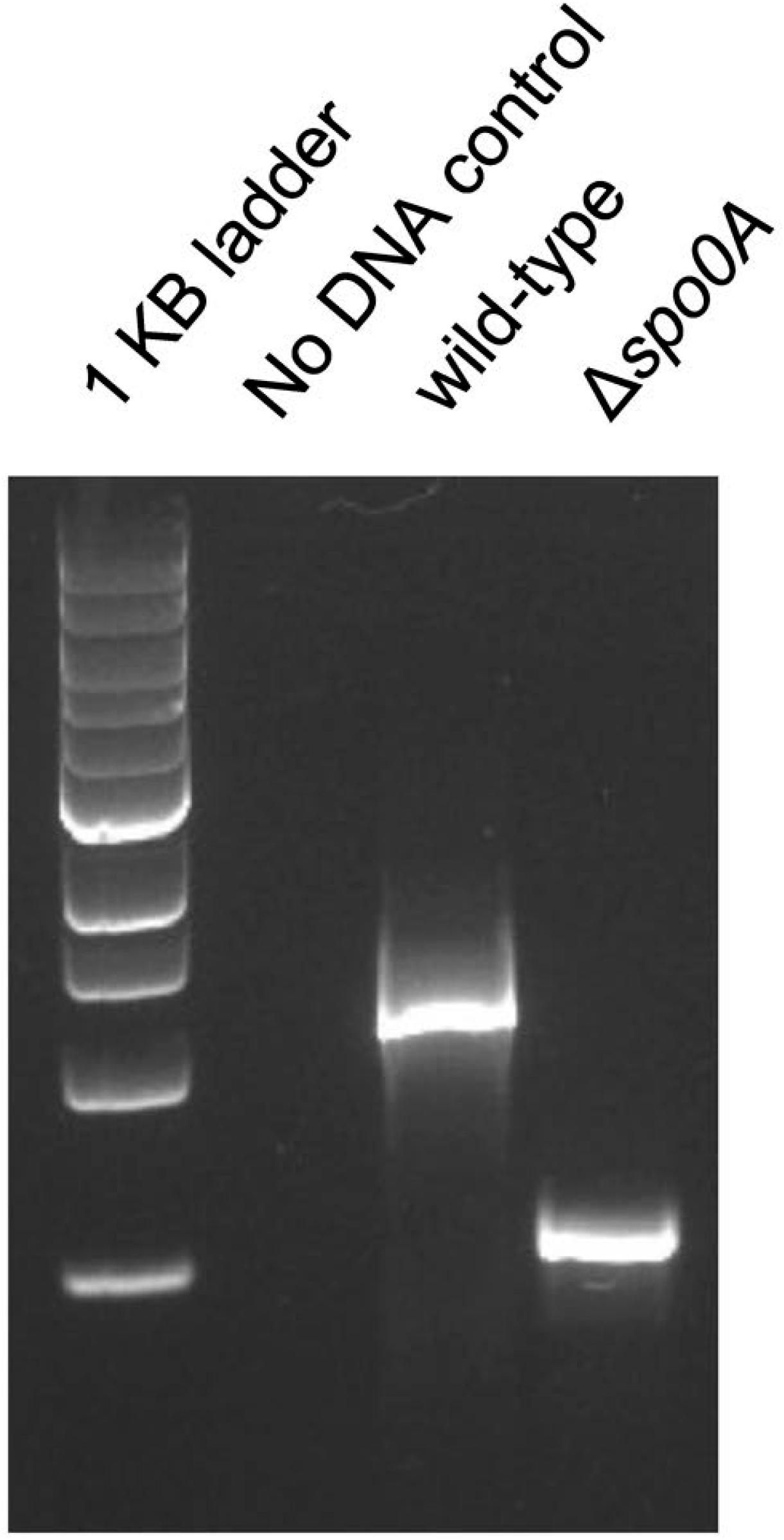
Deletion of *spo0A* in *C. difficile* R20291

Table S1: Complete list of gene expression fold-change from RNA-seq of wild-type and *selD* mutant strains.

Table S2: Strains and plasmids used in this study.

Table S3: Oligonucleotides used in this study.

